# Cortical Overgrowth in a Preclinical Forebrain Organoid Model of *CNTNAP2*-Associated Autism Spectrum Disorder

**DOI:** 10.1101/739391

**Authors:** Job O. de Jong, Ceyda Llapashtica, Kevin Strauss, Frank Provenzano, Yan Sun, Giuseppe P. Cortese, Karlla W. Brigatti, Barbara Corneo, Bianca Migliori, Steven A. Kushner, Christoph Kellendonk, Jonathan A. Javitch, Bin Xu, Sander Markx

**Affiliations:** Department of Psychiatry, Vagelos College of Physicians & Surgeons, Columbia University, New York, NY, USA; Division of Molecular Therapeutics, New York State Psychiatric Institute, New York, NY, USA; Department of Pharmacology, Vagelos College of Physicians & Surgeons, Columbia University, New York, NY, USA; Clinic for Special Children, Strasburg, Pennsylvania, USA; Taub Institute for Research on Alzheimer’s Disease and the Aging Brain, Department of Neurology, Columbia University, New York, NY, USA; Stem Cell Core Facility, Columbia University, New York, NY, USA; Department of Psychiatry, Erasmus MC University Medical Center, Rotterdam, The Netherlands; Tech4Health and Neuroscience Institutes, NYU Langone Health, New York, NY, USA

**Author notes:** Corresponding author: Sander Markx, M.D., Bin Xu, Ph.D.

## Abstract

Autism spectrum disorder (ASD) represents a major public health burden but translating promising treatment findings from preclinical non-human models of ASD to the clinic has remained challenging. The recent development of forebrain organoids generated from human induced pluripotent stem cells (hiPSCs) derived from subjects with brain disorders is a promising method to study human-specific neurobiology, and may facilitate the development of novel therapeutics.

In this study, we utilized forebrain organoids generated from hiPSCs derived from patients from the Old Order Amish community with a rare syndromic form of ASD, carrying a homozygous c.3709DelG mutation in *CNTNAP2* and healthy controls to investigate the effects of this mutation on cortical embryonic development.

Patients carrying the c.3709DelG mutation in *CNTNAP2* present with an increased head circumference and brain MRI reveals an increase in gray matter volume. Patient-derived organoids displayed an increase in total volume that was driven by an increased proliferation in neural progenitor cells, leading to an increase in the generation of cortical neuronal and non-neuronal cell types. The observed phenotypes were rescued after correction of the pathogenic mutation using CRISPR-Cas9. RNA sequencing revealed 339 genes differentially expressed between patient- and control-derived organoids of which a subset are implicated in cell proliferation and neurogenesis. Notably, these differentially expressed genes included previously identified ASD-associated genes and are enriched for genes in ASD-associated weighted gene co-expression networks.

This work provides a critical step towards understanding the role of *CNTNAP2* in human cortical development and has important mechanistic implications for ASD associated with brain overgrowth. This *CNTNAP2* organoid model provides opportunity for further mechanistic inquiry and development of new therapeutic strategies for ASD.

## INTRODUCTION

Autism spectrum disorder (ASD) and related neurodevelopmental disorders represent a major public health burden (1). To better understand these conditions, mechanistic insight into their development is needed. Rodent models with modified genes associated with ASD have facilitated this but translating promising preclinical findings to the clinic has remained enormously challenging (2). This is at least partly due to differences in the biology between species and underscores the need for elucidating human-specific disease mechanisms for psychiatric disorders. Forebrain organoids generated from human induced pluripotent stem cells (hiPSCs) that are derived from subjects with well-characterized brain disorders are a promising method to study human-specific neurobiology. Indeed, brain organoid modeling of ASD and related neurodevelopmental disorders may help reveal critical disease phenotypes and associated disease mechanisms that may not be recapitulated in animal models (3–6).

A homozygous *loss-of-function* (LoF) mutation, c.3709DelG, in contactin-associated-protein-like 2 *(CNTNAP2)* causes a rare and severe neurodevelopmental syndrome characterized by ASD, intellectual disability, early-onset epilepsy and an increase in head circumference – collectively termed Cortical Dysplasia Focal Epilepsy (CDFE) syndrome (7). The inherited mutation and associated syndrome were first described in a family from the genetically isolated Old-Older Amish population (7). Additional cases carrying different homozygous LoF mutations were subsequently discovered in subjects outside the Amish founder population, all presenting with a similar constellation of symptoms (8, 9). Thus far, no homozygous LoF mutations have been described in healthy control individuals, indicating that these are consistently associated with a severe and fully penetrant phenotype.

*CNTNAP2* encodes a cell adhesion molecule that is structurally related to the neurexins, but has shown to be functionally distinct, fulfilling roles in both peripheral and central nervous system development and physiology (10). Known functions of *CNTNAP2* have been delineated in a number of *Cntnap2* knock-out and knock-down rodent models, and include critical roles in potassium channel clustering in myelinated fibers (11) as well as dendritic arborization (12), synaptic function (13) and neuronal migration (14). Interestingly, *Cntnap2* null mice display several behavioral phenotypes reminiscent of the neurodevelopmental syndrome associated with homozygous LoF mutations, including deficits in social interaction and cognition, an increase in repetitive-stereotyped behaviors, and a lowered seizure threshold (14).

Little is known about the function of *CNTNAP2* in the human brain. Human expression data shows high abundance in different cortical areas during embryonic development (15). Histopathology and MRI findings from patients from the Old-Order Amish population and diagnosed with CDFE-syndrome show cortical dysplasia, cortical grey matter thickening, ectopic neurons in the white matter tracts and increased hippocampal cell number (7). Together, these findings imply an early embryonic origin of pathophysiology. As embryonic development is most faithfully recapitulated with brain organoid modeling systems as shown by transcriptome analyses (16, 17), we generated brain organoids to study how disruption of *CNTNAP2* affects cortical embryonic development.

To this end, we generated hiPSC lines from subjects from the Old-Order Amish founder population, all of whom had been diagnosed with CDFE-syndrome and carried the homozygous c.3709DelG mutation in *CNTNAP2.* Using these lines we generated forebrain organoids, and compared these to organoids generated from healthy controls from the same population to examine the effects of the mutation on cortical development. We show an increase in volume of patient-derived organoids compared to controls. This enlargement of organoid volume results from increased proliferation of neural progenitor cells (NPCs) which, in turn, leads to an increased generation of both cortical neurons and non-neuronal cell types. RNA sequencing revealed 339 differentially expressed genes, including genes involved in cell proliferation and neurogenesis. Finally, we show that a portion of these differentially expressed genes have also been associated with idiopathic and syndromic autism and are also present in autism-associated gene co-expression networks. Together, these findings provide a possible explanation for the increased head circumference and cortical gray matter thickening that has been observed clinically in these patients (Strauss, 2006). Our work provides a critical step towards understanding the role of *CNTNAP2* in brain development. The increase in NPC proliferation observed in patient-derived organoids provides a disease mechanism that could be further investigated for developing new treatments of *CNTNAP2*-associated and other forms of ASDs.

## METHODS

### Ascertainment of Subjects and Clinical Phenotypes

The study was approved by the Lancaster General Hospital (LGH) Institutional Review Board (IRB) (LGH IRB protocol number 2008-095). Parents of all subjects provided written informed consent to participate on behalf of their children. Neuropsychiatric diagnoses and head circumferences measurements were done during routine clinical visits for pediatric care at the Clinic for Special Children in Strasburg, Pennsylvania. All work with the hiPS cell lines and derived neurons and forebrain organoids took place under a New York State Psychiatric Institute (NYSPI) IRB-approved protocol (NYSPI IRB protocol number 7500).

### MRI analysis

We quantified total gray matter volume relative to the total brain volume of 5 whole brain MRIs from patients carrying the c.3709DelG mutation in *CNTNAP2*, and compared these to 4 ‘template MRIs’ that were generated from 10 combined whole-brain MRIs of age-matched healthy children. We used Statistical Parametric Mapping for image segmentation and summed gray matter and white matter masks to calculate total brain volume. For each image, the sum value of voxels for each tissue class was used to generate a ratio between gray matter and white matter.

### hiPSC generation and characterization

hiPSC lines were generated from three patients with *CNTNAP2*-associated ASD carrying the homozygous c.3709DelG mutation and three healthy controls from the same Amish population who did not carry mutation, by a non-integrating Sendai virus-based reprogramming method as previously described (18). Karyotyping was performed on twenty G-banded metaphase cells at 450–500 band resolution as previously described (19). All hiPSC lines used in this study were between passage 10 and 25. The difference in passage number for each pair of patient- and control-derived line never exceeded 3 passages.

### Forebrain organoid culture

Forebrain organoids were generated from patient- and control hiPSC-lines using a previously published organoid differentiation protocol (20), with minor modifications. Briefly, hiPSCs were dissociated with 0.5 mM EDTA in PBS and triturated to generate a single cell suspension. A total of 10,000 cells were then plated into each well of an ultra-low-attachment 96-well plate (Nunc) to form embryoid bodies (EBs) in medium containing mTesR1, 1 µg/ml heparin and Penicillin-Streptomycin antibiotics. EBs were exposed to medium containing the ROCK inhibitor Y27632 (50 µM) for the first 24 hrs, followed by 5 days without interference in 96-well plates. On day 5, the medium was switched to medium containing DMEM/F12 (1:1) (Gibco; 11330), N-2 supplement (Gibco; 17502-048), MEM-NEAA (Invitrogen), Glutamax (Invitrogen 35050-061), Penicillin-Streptomycin and 0.1 mM β-Mercaptoethanol, Dorsomorphin (2 µM), SB431542 (10 µM) and IWR1e (3 µM). On day 25, 0.2% Chemically Defined Lipid Concentrate was added to this medium until day 39. On day 40, EBs were transferred to ultra-low attachment 24 well plates with 1 ml of medium per organoid/well. Between day 40 and 80, 1 ml of medium containing DMEM/F12 (1:1) (N-2 supplement MEM-NEAA, Glutamax, Penicillin-Streptomycin and 0.1 mM β-Mercaptoethanol, 10% fetal bovine serum (FBS) and 1% Matrigel (Corning) was replaced weekly. After day 80 medium was replaced once per week. Matrigel was added bi-weekly to prevent excess Matrigel deposition on the outer shell of the organoid.

### Immunohistochemistry

See table S1 for all antibodies and respective concentrations used. Organoids were fixed for 4 h in 10% formalin at room temperature, cryo-protected in 15% sucrose, dissolved in PBS for 4 h, and then placed in 30% sucrose in PBS overnight. The next day organoids were embedded in optimal cutting temperature (O.C.T.) solution and cut into 14 μm sections on a Leica Cryostat. After heat-induced epitope retrieval in 40 mM Sodium Citrate/ 0.05% Tween-20 (pH6), tissue sections were blocked for 1h in 10% horse serum in PBS containing 0.1% TritionX-100 (PBSTx0.1). Primary antibodies were incubated overnight at 4 °C in PBSTx0.1. Secondary antibodies were incubated for 1h at room temperature and nuclei were counterstained with 300 nM DAPI solution. Sections were mounted on glass slides using Prolong antifade solution and imaged on a LEICA SP8 confocal microscope.

### Electrophysiology

Measurement of Electrophysiology Properties. Whole-cell patch-clamp recordings were performed as previously described (21). EBs were dissociated into single cells after rosettes were selected, plated onto glass coverslips coated with poly-L-ornithine (50µg/ml) and laminin (5µg/ml), and maintained *in vitro* for 60 days. The external solution was (in mM): 145 NaCl, 5 KCl, 2 CaCl_2_, 2 MgCl_2_, 10 glucose, and 10 HEPES (pH 7.3 using NaOH and the osmolarity adjusted to 325 mOsm with sucrose). Patch pipettes (open tip resistance 5–8 MO) were filled with a solution containing (in mM): 130 CH_3_KO_3_S, 10 CH_3_NaO_3_S, 1 CaCl_2_, 10 EGTA, 10 HEPES, 5 MgATP, and 0.5 Na_2_GTP (pH 7.3, 305 mOsm). Cells were voltage-clamped at −70 mV and resting membrane potential was measured immediately following the establishment of the whole-cell configuration. Membrane resistance and capacitance were calculated from the membrane potential changes in response to 1 s duration hyperpolarizing current steps increasing by 5 pA increments. Action potentials were evoked using this same current-step protocol, and were defined as a transient depolarization of the membrane which had a minimum rise rate >10 mV/ms and reached a peak amplitude > 0 mV. Intrinsic properties were measured from the first action potential. Duration was calculated from the full width at the half maximum voltage, with amplitude measured from the threshold potential to the maximum potential. Quantification was done using Igor Pro v.6 (Wavemetrics). Statistical comparisons were made using unpaired Student’s t test in GraphPad Prism v.7.

### Light-sheet microscopy

Whole, fixed organoids were stained in 300 nM DAPI in PBS for 30 mins to ensure staining of the nuclei of the outer shell of the organoid. Light sheet microscopy images were acquired on a Zeiss Z-1 Light sheet microscope from 6 angles in 45° degree increments starting at 0°.

### CRISPR-Cas9 Rescue-line generation

We randomly selected one of the three patient hiPSC lines for CRISPR genome editing.

*gRNA and donor template design.* Guide-RNAs were designed using *CT-finder* software (22). gRNAs were designed around the c.3709DelG mutation site in the *CNTNAP2* locus on chromosome 7q35. gRNA target A on the minus strand is 5’**-** GGGTCGGTGGCGGACGACATGGG-3’ and gRNA target B on the plus strand is 5’-GCACCTGGATCACCTGATTCAGG-3’. The donor template was a single stranded 181-bp oligonucleotide (see Table S2). This sequence is complementary to the antisense DNA strand and contains the wildtype DNA sequence and a silent blocking mutation near the 5’-end of gRNA target A. The donor template flanks the mutation site with 90bp on each side (Fig S2A-).

*hiPSC gene-editing using CRISPR Cas9*. We followed a previously described protocol (23) with minor modifications. Briefly, gRNA oligonucleotides were cloned into the plasmid pSpCas9n(BB)-2A-GFP encoding Cas9-nickase. ihPSCs were maintained in MTesR1 medium. Cells were passaged using 0.05 mM EDTA in PBS or Accutase. Prior to nucleofection, cells were treated with ROCK-inhibitor for 24 hours. Plasmids and donor template were introduced into the hiPSCs through nucleofection using Amaxa nucleofector II. 72 hours post nucleofection, GFP + cells were selected using FACS and grown as single cell-derived colonies onto irradiated MEF coated 35 mm dishes. Colonies with undifferentiated hiPSC morphology were manually picked and further grown on 96-well plates. Clones were genotyped using Sanger sequencing on PCR amplicons (see table S2 for primers).

### Viral transduction

pAAV.hSyn.eGFP.WPRE.bGH was a gift from James M. Wilson (Addgene plasmid # 105539; http://n2t.net/addgene:105539; RRID:Addgene_105539). The virus was added to the organoid growth media for 1 week at a concentration of 10^7^ virus particles per ml. Transduction efficiency was evaluated 10 days post-transduction using an epi-fluorescence microscope.

### Isotropic fractionation

We adapted a previously developed protocol for to accurately calculate total cell number and relative cell type fractions based on nuclear immunostaining (24). Briefly, organoids were fixed for 4h in 10% formalin at room temperature. After blocking for 4h in 10% horse serum in PBS containing 0.2% Triton X-100 (PBSTx0.2), primary antibodies were incubated PBSTx0.2 on whole organoids for 20h on a shaker at room temperature to ensure complete tissue penetration. Organoids were washed for 4 h in PBSTx0.2. Secondary antibodies were incubated for 20h in in PBS PBSTx0.2, followed by a 4 h wash in PBSTx0.2. Organoids were fixed for a second time for 20h, and then homogenized using a glass tissue homogenizer in 10 mM sodium citrate buffer containing 1% Triton X-100. Total cell counts were performed by counting DAPI-stained nuclei in a hemocytometer on an upright Zeiss epifluorescence microscope. Relative cell type fractions were calculated from images acquired on a Leica SP8 confocal microscope.

### BrdU labeling

Organoids were incubated using 10 µM BrdU dissolved in growth medium for 2 h at 37 °C, then washed twice with PBS and subsequently fixed for 4 h in 10% formalin at room temperature. We subsequently followed our immunostaining protocol to stain for BrdU after heat denaturing the DNA in a 40 mM sodium citrate buffer containing 0.05% Tween-20. For the BrdU/Ki67 quantification analysis, serial sections of 14 μm were made every fifth section, with 3-4 sections for each organoid.

### Total RNA isolation and RNA sequencing

Ten forebrain organoids per line (3 control-derived lines, 3 patient-derived lines) were collected at 8 weeks *in vitro*. Total RNA was isolated from hiPSC-derived cortical neurons using miRNeasy kit (Qiagen, USA) according to instructions of the manufacturer. RNA was suspended in RNase-free water. The concentration and purity of each sample was determined by spectrophotometer (ND1000; Nanodrop) and confirmed by Microchip Gel Electrophoresis (Agilent) using Agilent 2100 Bioanalyzer Chip according to the instructions of the manufactures. A poly-A pull-down step was performed to enrich mRNAs from total RNA samples (200 ng to 1 µg per sample, RIN > 8 required) and proceeded on library preparation by using Illumina TruSeq RNA prep kit. Libraries were then sequenced using Illumina HiSeq2000 at the Columbia Genome Center. Multiplex samples with unique barcodes were mixed in each lane, which yields targeted number of single-end 100 bp reads for each sample, as a fraction of 180 million reads for the whole lane. RTA software (Illumina) was used for base calling, and bcl2fastq (version 1.8.4) for converting BCL to fastq format, coupled with adaptor trimming. The reads were mapped to a reference human genome (hg19) using Tophat20 (version 2.0.4) with four mismatches (--read-mismatches = 4) and 10 maximum multiple hits (--max-multihits = 10).

### Differential expression analysis

DESeq2, an R package based on a negative binomial distribution that models the number reads from RNA-seq experiments and tests for differential expression, was used to determine differentially expressed genes (DEGs) between mutants and control samples. The list of significantly DEGs was defined at false discovery rate (FDR) adjusted p-value (padj) < 0.05.

### Gene ontology analysis

To determine a common functional relationship among the top DEGs, the enrichment of biological processes was tested using Metascape (25) with default settings. Gene lists of upregulated and downregulated DEGs were analyzed separately.

### Permutation tests

Permutation tests were performed by generating 10000 gene lists of equal size to the differentially expressed genes. We then compared these randomly generated gene lists to the gene lists of interest (WGCNA networks, ASD-SFARI genes) as well as control gene lists derived from other common diseases, and tested whether the overlap between DEGs and permuted gene lists differed in a statistically significant manner from the randomly generated gene lists.

### Image analyses

Confocal and light sheet microscopy images were processed using ImageJ software. BrdU/Ki67 quantifications and nuclear fraction quantifications were carried out using Cell Profiler software.

### Statistical analyses

All statistical analyses were performed using *R* programming language. The graphs in the manuscript display boxplots showing median, first, and third quartiles. Whiskers show the largest and smallest values no further than 1.5 times the interquartile range from the first and third quartile, respectively. Data points outside of this range are plotted individually. Comparisons between patient and control-derived organoids were done using a linear mixed model with genotype status designated as a fixed effect and cell line as a random effect. Relationships between two continuous variables were tested using linear regression. Statistical significance from the mixed model was calculated using Likelihood Ratio Tests (LRT). For comparisons of the CRISPR rescue-line to its original patient line, and the gray matter volume analysis, we used student’s t-tests. P-values < 0.05 were considered statistically significant. Outliers were never removed from any of the analyses.

## RESULTS

### Clinical description of patients carrying c.3709DelG mutation in *CNTNAP2*

Because the sample size of the initial cohort in which the increased head circumference was described was small (7), we expanded this assessment by measuring the head circumferences of 17 male and 20 female pediatric patients from the Old-Order Amish founder population all carrying the inherited homozygous c.3709DelG mutation in *CNTNAP2* and all presenting with the same syndromic ASD. We compared these data to head circumference reference data published by the WHO (26). This analysis revealed an increased head circumference in both male and female patients compared to the WHO mean (weighted z-score = 2.93, p <0.003 (Fig S1A). To expand upon the initial description of increased gray matter thickness (7), we analyzed brain MRIs from 5 patients carrying the c.3709DelG mutation in *CNTNAP2* and compared these to age-matched clinical templates. We found an increase in total gray matter (GM) volume relative to total brain volume (TBV) in patients compared to controls (Patients: GM/TBV= 0.73; Controls: GM/TBV= 0.54; t-test, p = 0.004, n= 5 patients, 4 control templates (Fig. S1B).

### hiPSC and Forebrain Organoid Generation and Characterization

We generated hiPSC lines from monocytes of three human female patients (ages 10, 12 and 17 years) carrying the homozygous c.3709DelG mutation in *CNTNAP2* (Fig. 1A) and three sex-matched healthy control subjects from the same population. All 3 patients were assessed by a pediatrician (KS) and were diagnosed with epilepsy and a syndromic form of ASD, characterized by cognitive-, behavioral-, and language regression and delayed developmental milestones starting around age 1.

**Figure 1.**
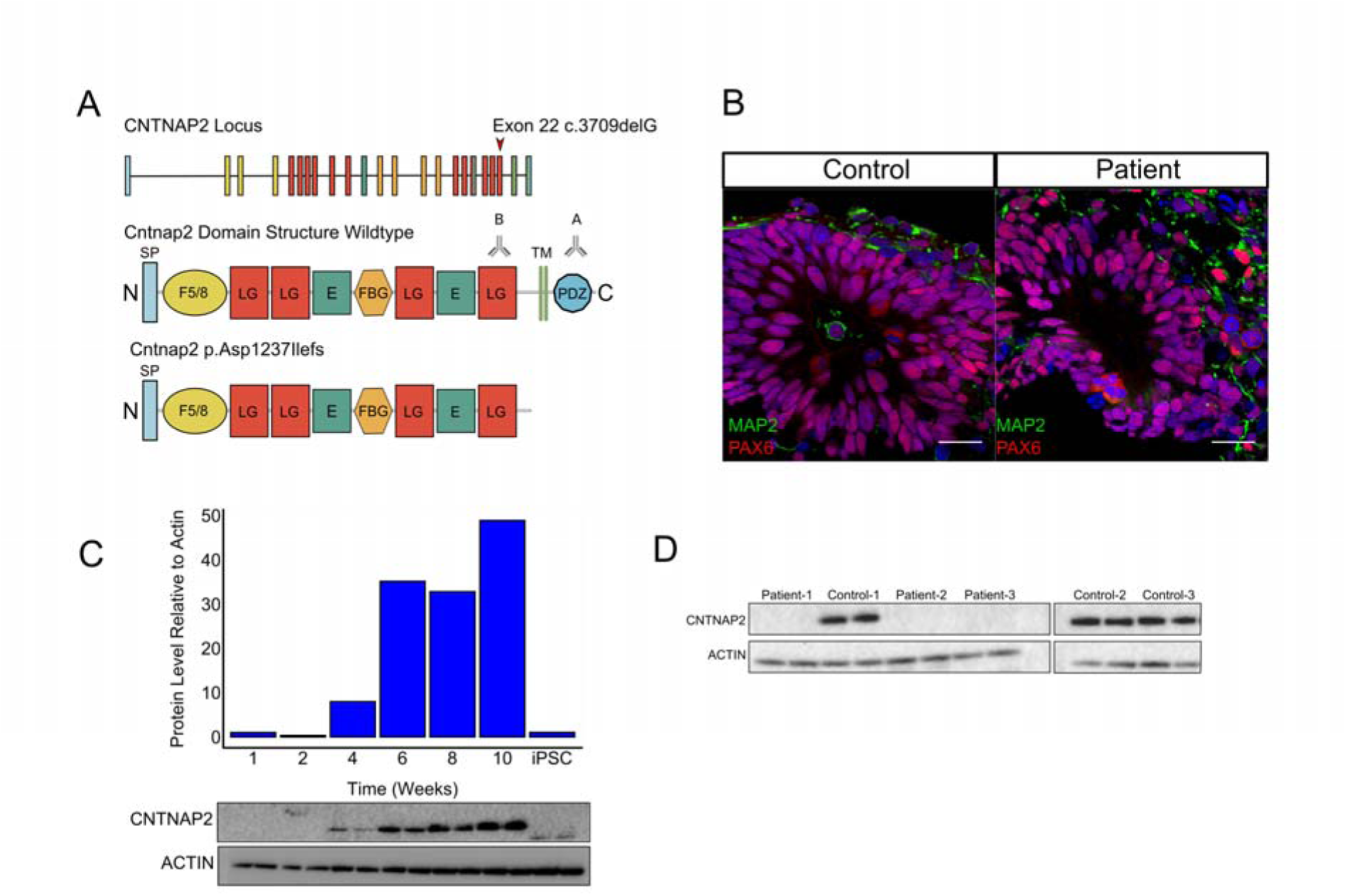
CNTNAP2 expression and forebrain organoid characterization. A. *CNTNAP2* locus and protein domain structures of wildtype and mutant protein. The 24 exons of *CNTNAP2* are color-coded based on domain structures of the full-length protein. Deletion of a guanine base at position 3709 in exon 22 (indicated with red arrow) of the gene results in a substitution of aspartic acid for isoleucine at amino acid position 1237, leading to a shift in reading frame and a stop codon at amino acid position 1253. The truncated variant of the protein (p.Asp1237Ilefs) is terminated right before the C-terminal transmembrane domain. Antibody A (intracellular) is a polyclonal antibody directed against the C-terminal end of the protein. Antibody B (extracellular) is a monoclonal antibody directed against the Laminin-G domain upstream of the transmembrane domain (also see table S1 for antibodies). SP= signal peptide; F5/8= discoidin/neuropilin homology domain; LG= Laminin G domain; E= EGF-Like Domain; FBG= Fibrinogen-Like Region; PDZ = PDZ-interaction domain. B. Both control-derived and patient-derived organoids show PAX6-expressing neural progenitor cells in ventricular zone-like regions after 4 weeks in culture. Neuronal dendrosomatic marker MAP2-expressing cells are preferentially located in the outer zones of the cortical organoids. Scale bar represents 50 μm. C. Western blot analysis showing increasing *CNTNAP2* expression levels over time relative to actin in a control-derived organoid line, roughly corresponding to the developmental pattern of *CNTNAP2* expression pattern in human embryonic brain. D. Western blot analysis showing presence of full-length *CNTNAP2* protein levels in control-derived organoids and absence in patient-derived forebrain organoids using an antibody directed against the C-terminal end of the protein (Antibody A in panel A).

To confirm stemness in hiPSC lines used for organoid generation we performed RT-PCR for markers NANOG and OCT4 (Fig. S1C), we performed G-band karyotype analysis to ensure the abcense of chromosomal abnormalities in all patient and control derived cell lines (Fig. S1D We confirmed the genotypes of patient- and control-derived hiPSCs using Sanger sequencing (Fig. S1E).

The forebrain organoids generated from both patients and healthy controls displayed similar characteristics of early telencephalic brain development. After 4 weeks in culture, both patient- and control-derived organoids showed ventricle-like structures composed of NPCs expressing forebrain-specific transcription factor PAX6, (Fig. 1B). Dorsal forebrain identity was further confirmed by immunostaining for transcription factor FOXG1 (Fig. S1F). Organoids from both genotype groups demonstrate a clear pattern of MAP2-expressing neurons positioned around the ventricle-like structures and preferentially located in the outer layers of the organoids, indicative of initial NPC differentiation and subsequent migration towards the outer cortex-like region (Fig. 1B). After 8 weeks in culture, neurons that were dissociated from both patients and healthy controls for whole-cell patch-clamp recording displayed a consistent pattern of mature electrophysiological characteristics (Fig. S1G and H) indicating that the organoid differentiation protocol was implemented successfully.

### *CNTNAP2* expression in forebrain organoids

To determine the *CNTNAP2* expression pattern control-derived organoids, we performed immunoblotting, which revealed expression starting around 4 weeks in culture that increased over time (Fig. 1C). Immunoblotting for CNTNAP2 utilizing an antibody (Antibody A, intracellular) Fig. 1A) directed against the c-terminus that is deleted in the c.3079delG mutation showed that full-length CNTNAP2 protein could not be detected in patient-derived organoids while it was indeed detected in control-derived organoids (Fig. 1D). We then used an antibody directed against the laminin G domain of the protein located upstream of the transmembrane domain (Antibody B Fig. 1A), which is predicted to be present in the truncated CNTNAP2 protein. Notably, while immunoblotting with antibody B revealed *CNTNAP2* expression in the control samples, no protein was detected in patient-derived organoids suggesting that the truncated *CNTNAP2* mRNA and/or protein are unstable (Fig. S1I).

To compare these expression studies in organoids to expression in the intact brain, we analyzed data from the publicly available gene-expression database BrainSpan (27), which characterizes the embryonic spatiotemporal expression pattern of in the developing human brain. These data show the highest *CNTNAP2* expression levels in the first trimester of embryonic development in multiple cortical areas, including orbitofrontal, prefrontal and temporal cortices (Fig. S1J). Other areas of high gene expression during embryonic development include the basal ganglia and cerebellum (Fig. S1I). The BrainSpan data are in line with previous expression studies on CNTNAP2 in the human embryonic brain (15, 28). These results indicate that the *CNTNAP2* expression pattern in organoids are in concordance with *in vivo* gene expression, and thus indicate that forebrain organoids serve as a well suited preclinical neuronal model system to study the effects of this homozygous LoF mutation on early human cortical development.

### Patient-derived forebrain organoids display an increase in volume

To investigate whether the brain overgrowth characteristic of these patients could be recapitulated in our model system, we set out to assess the size of patient- and control-derived organoids over time by measuring the projected surface area of the organoid using 2D bright field microscopy images (Fig. 2A). At 13 weeks *in vitro*, patient-derived organoids generated from an equal number of iPCSs and comparable in size at 4 weeks around the onset of *CNTNAP2* expression, showed a 1.5-fold increase in 2D projected surface area compared to control-derived organoids (LRT, p = 0.002, n= 6-7 organoids/line) (Fig 2B).

**Figure 2.**
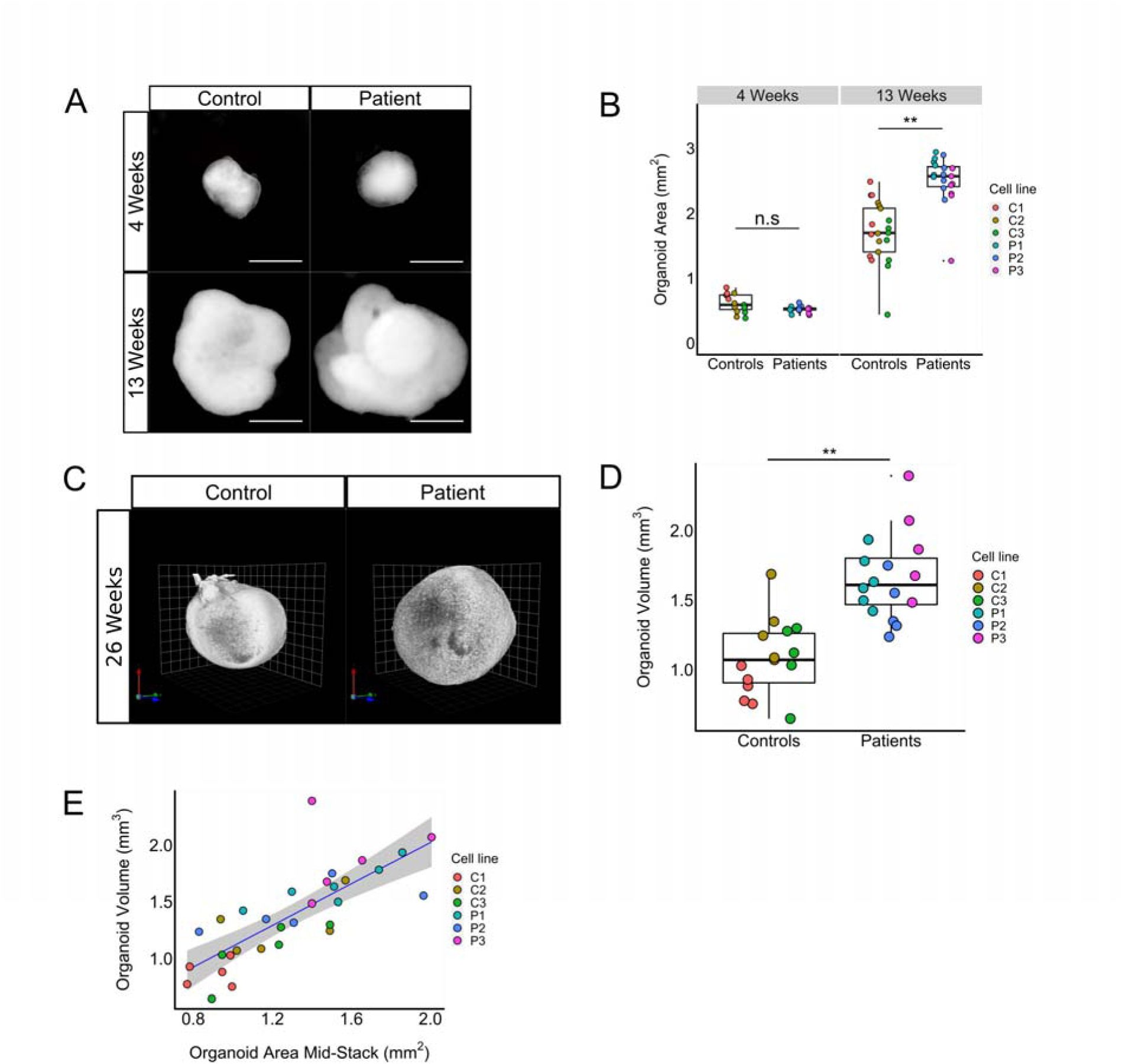
Patient-derived forebrain organoids display an increase in volume. **(A and B).** Representative image of bright field images of patient- and control-derived organoids over time (A) and quantifications based on 2D bright field image measurements (B), showing a 1.5-fold increase in area of patient-derived organoids versus control-derived organoids at week 13 (LRT, p = 0.002, χ2(1)= 9.42), while no difference in size was observed at week 4. Scale bar represents 500 μm. **(C and D).** Representative image of light-sheet microscopy images of control- and patient-derived cortical organoids at 26 weeks *in vitro* (C) and volumetric quantification of light-sheet images showing a 1.5-fold volumetric increase for patient-derived cortical organoids compared to controls (D); LRT, p = 0.005, χ2(1)= 7.85, n=xx). **E.** Scatterplot showing the relationship between the area of a middle z-stack of a light-sheet image of a forebrain organoid and the volume at 26 weeks as measured by the 3D rendering of the light sheet image. We constructed a linear model of volume as a function of the area of the middle z-stack (blue line) (F(1,29)= 44.59, R^2^ = 0.606, p = 2.529e-07). Shaded in grey is the 95% confidence interval estimated for the linear model. This result indicates that 2D bright field microscope images can be used to estimate organoid volume. Boxplots display median, first and third quartiles, and whiskers showing the largest and smallest values no further than 1.5 times the inter quartile range from first and third quartile, respectively. Data points outside of this range are plotted individually. * = p < 0.05; ** = p < 0.01; *** = p < 0.001

As the organoids are not spherical but irregularly shaped, we used light sheet microscopy to more precisely quantify organoid volume (Fig. 2C). We found a 1.5-fold volumetric increase for patient-derived organoids compared to controls (LRT, p = 0.005) (Fig. 2D). We then utilized light-sheet imaging to investigate whether the total surface area of the middle z-section could be used as an reliable estimate for organoid volume using a linear model, which was indeed the case (Linear model, R^2^ = 0.606, p < 0.001) (Fig. S2A). We therefore proceeded using 2D brightfield microscopy images for quantifying organoid volume.

### Increased organoid volume is driven by an increased total cell number caused by increased NPC proliferation

To establish the factors that contribute to the volumetric increase of the patient-derived organoids, we utilized isotropic fractionation (24) to estimate the total cell number in the organoids. We found a 2.5-fold increase in total cell number after 6 weeks in culture for patient-derived organoids compared to controls (LRT, p <0.001, n =4 organoids/line) (Fig. 3A). As expected, organoid surface area and total cell number correlated positively with each other at both 6- and 13 weeks *in vitro* (Fig. 3B and Fig. S3A). To determine whether cell size also contributes to the volumetric increase of the organoids, we measured the neuronal soma of AAV-Synapsin-GFP transduced organoids, and found no difference in neuronal soma volume between genotype groups (LRT, p= 0.69) (Fig S3B). We then estimated the cell cycle length by co-labeling with BrdU and Ki67, and found that proliferating cells have a shorter cell cycle (Average cell cycle time (Tc) = 54h) for patient-derived organoids compared to controls (Tc= 83h) (LRT, p =0.02, 1.54-fold change = 1.54, n = 3 organoids/line) (Fig. 3C and D). Co-labeling of BrdU with PAX6 confirmed that most proliferation took place in the PAX6+ NPCs (Fig. S3C) in organoids from both genotype groups (LRT, p=0.46, n= 3 organoids/genotype). Next, we quantified the size of the progenitor pool by calculating the absolute numbers of PAX6+ cells at 6 weeks *in vitro*. We found a 2.6-fold increase in absolute numbers of NPCs in the patient-derived organoids compared to controls (LRT, p <0.001, n=4 organoids/line) (Fig. 3EFG). These data show that increased proliferation activity of NPCs leads to an increase in the progenitor pool, which is a major contributor to the volumetric increase observed in patient-derived organoids.

**Figure 3.**
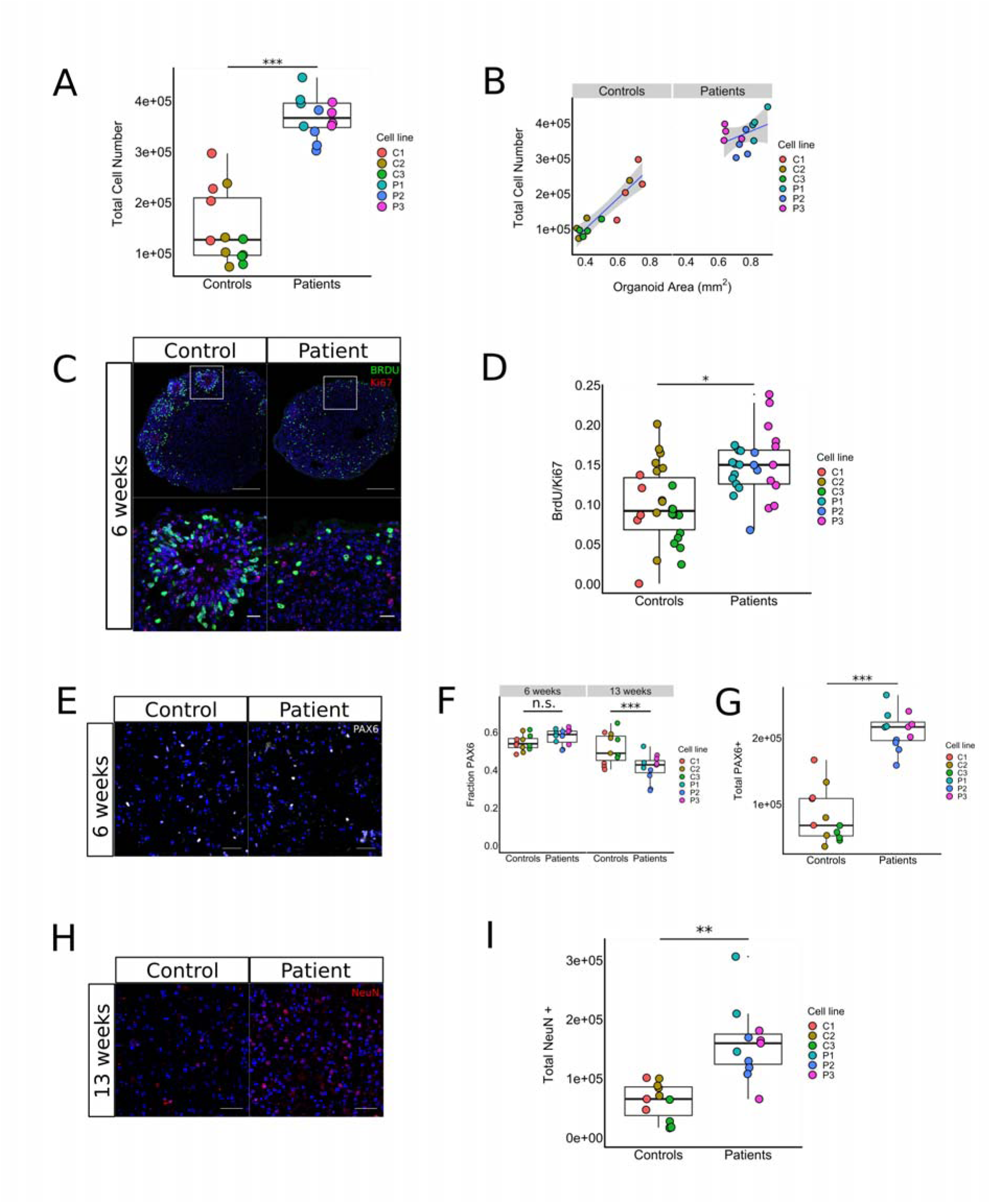
Volumetric increase is driven by increased NPC proliferation, which leads to expansion of progenitor pool and increased cortical neurogenesis. **(A and B).** Total cell number quantification of patient- and control-derived organoids after 6 weeks in culture, showing a 2.5-fold increase in total cell number for patient-derived organoids (LRT, p = 0.0003, χ2(1)= 13.25) (A), and the linear relationship between 2D organoid surface area and total cell number for patient- and control-derived organoids after 6 weeks in culture. Controls: F(1,10) = 50.22, R^2^= 0.83, p <0.001; patients: F(1,10) = 1.82, R^2^= 0.15, p = 0.21. **(C and D).** Representative image of organoid sections stained for BrdU and Ki67 after 6 weeks in culture (C) and quantification (D) of ratio BrdU/Ki67 between patient- and control-derived organoids, showing a 1.5-fold increase in BrdU/Ki67 ratio for patient-derived organoids (LRT, p = 0.02, χ2(1)= 5.55). Using the formula Tc = Ts/(BrdU+/Ki67+), where Tc = cell cycle length, and Ts = S-phase length, estimated mean cell cycle lengths for patient- and control-derived organoids are ∼ 54 and ∼ 83 hours, respectively. Scale bar represents 50 μm. **(E, F and G).** Representative image of confocal microscopy images of nuclei from homogenized control and patient-derived organoids cultured for 6 weeks and stained for NPC-marker PAX6 (E), quantification of fractions of PAX6 + cells at 6 and 13 weeks *in vitro* (F) and total number of PAX6+ cells at 6 weeks *in vitro* (G), showing a 2.6-fold increase in total PAX6+ cells at 6 weeks (LRT, p = 0.0003, χ2(1)= 13.35), with an equal fraction of PAX6* cells at six weeks, followed by a 1.2-fold decrease in PAX6 fraction at 13 weeks (LRT, p=0.0008, χ2(1)=11.23,) in support of increased differentiation towards a non-NPC cell fate. **(H and I).** Representative image of confocal microscopy images of nuclei from homogenized patient- and control-derived organoids cultured for 13 weeks and stained for post mitotic neuronal marker NeuN (H), and quantification of total amount of neurons generated at 13 weeks (I), showing a 2.6-fold increase for patient-derived organoids compared to controls (LRT, p=0.006, χ2(1)=7.57). Boxplots display median, first and third quartiles, and whiskers showing the largest and smallest values no further than 1.5 times the inter quartile range from first and third quartile, respectively. Data points outside of this range are plotted individually. * = p < 0.05; ** = p < 0.01; *** = p < 0.001

### Patient-derived cortical forebrain organoids display increased neurogenesis

To further characterize the nature of the overgrowth phenotype we investigated whether in addition to cycling at a higher speed, NPCs from patient-derived organoids also displayed abnormal cell cycle exit properties. Both measures have been shown to be altered in a brain organoid KO model of PTEN, an syndromic ASD gene (29). Ki67 labeling revealed that at 6 weeks *in vitro*, a lower fraction of cells were immunopositive for Ki67 in patient-derived organoids compared to controls, suggesting a faster differentiation towards non-progenitor cell fates that follows initial progenitor pool expansion, although this finding did not reach the threshold for statistical significance. (LRT, p =0.07, fold-change = 0.85, n=3 organoids/line) (Fig. S3D). To test this hypothesis further, we quantified the fraction of PAX6+ cells at 13 weeks *in vitro* and found a 1.2-fold decrease in the fraction of NPCs in patient-derived organoids compared to controls, which is consistent with the increase in the differentiation speed of NPCs (LRT, p< 0.001) (Fig. 3F).

We then investigated whether the increase in NPC proliferation and differentiation rate leads to a corresponding increase in the number of cortical neurons, which has been previously observed in post-mortem brains from patients diagnosed with ASD having increased head circumference (30). To do this, we quantified the relative and absolute numbers of post-mitotic neurons generated in the organoids by immunostaining nuclei for NeuN. We found a 2.6-fold increase in the absolute number of neurons generated (LRT, p = 0.006) (Fig. 3H and I) in patient-derived organoids, while NeuN fractions did not differ across genotype groups (LRT, p=0.66) (Fig. S3E). Consistent with this, the number of non-neuronal cell types was also increased (LRT, p= 0.004, 2.23-fold change, n = 3-4 organoids/line) (Fig. S3F). This finding further supports increased proliferation as the major driver of the observed increase in total number of neuronal cells generated in the patient-derived organoids.

### Site-specific repair of c.3709DelG mutation using CRISPR-Cas9 rescues cortical overgrowth phenotypes

To confirm the causal relationship between the homozygous *CNTNAP2* c.3709DelG mutation and the cortical overgrowth disease phenotypes, we used clustered regularly interspaced short palindromic repeats (CRISPR)-Cas9 to generate an isogenic ‘rescue’ line by reintroducing the guanine base at position 3709 in one of the patient-derived hiPSC lines (Fig. 4A). Sanger sequencing confirmed the wildtype DNA sequence (Fig. 4A). Immunoblotting confirmed the presence of the CNTNAP2 protein in the organoids derived from the CRISPR-line (Fig. 4B). We subsequently repeated the experiments by following the same organoid differentiation protocol described above. Organoids generated from the CRISPR-rescue hiPSC line had a 1.5-fold reduction in 2D projected surface area compared to the organoids derived from the unedited patient line (Fig. 4C-D) (t-test, p = 0.0014) – which corresponds with the observed baseline difference in 2D surface area between patient- and control-derived organoids. In addition, all other overgrowth phenotypes, including increased total cell number (t-test, p =0.007) (Fig. 4E), increased PAX6+ progenitor pool at 6 weeks (t-test, p=0.002) (Fig. 4F), decreased number of neurons (t-test, p =0.008) (Fig. 4G) and decreased fraction of PAX6 + cells at 13 weeks (t-test, p=0.04) (Fig. 4H) – were rescued to comparable numbers observed in the organoids generated from healthy controls.

**Figure 4.**
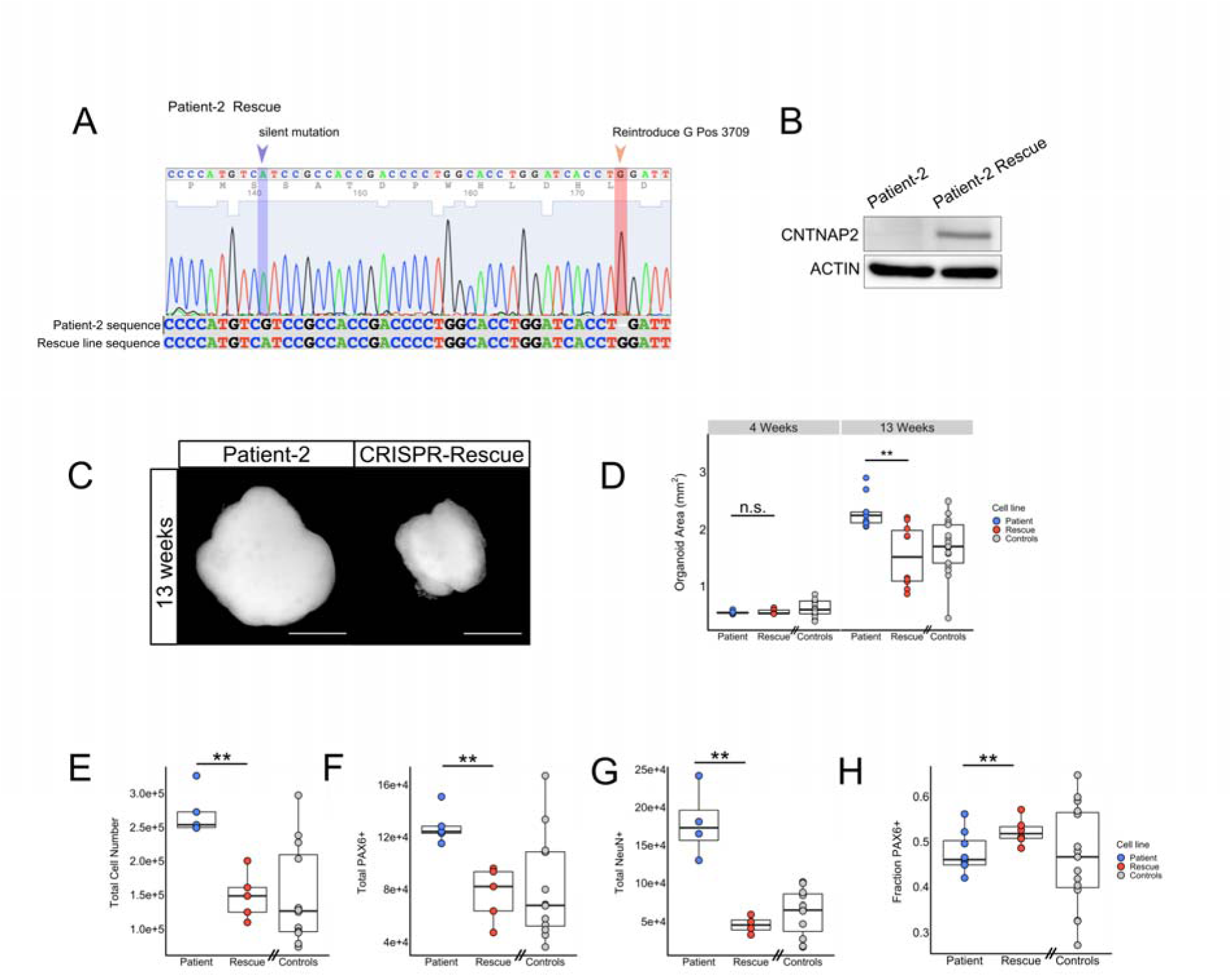
CRISPR-mediated repair of pathogenic c.3709DelG mutation rescues overgrowth phenotypes. **(A and B)** Sanger sequencing trace file of iPSC-line derived from patient-2 after CRISPR-Cas9 genome editing. The rescued line shows a guanine base introduced at cDNA position 3709 and a silent blocking mutation at position 3675. Sequence is aligned to original patient-2 DNA sequence (A). Immunoblotting confirms the presence of *CNTNAP2* protein in organoids generated from the CRISPR-rescued line, and absence of the protein in the parental patient-2 cell line-derived organoids (B). **(C and D).** Representative image of bright field microscopy images of organoids derived from patient-2 and CRISPR-rescued line at 13 weeks *in vitro* (C), and corresponding quantification showing a 1.5-fold decrease in area for the CRISPR rescue line compared to the unedited patient line (D) (T-test, T(14.1) = 3.93, p = 0.0014). Grey dots represent pooled control data points also displayed in figure 2A. Scale bar represents 500 μm. **(E, F and G).** Quantifications of total cell number at 13 weeks (E), total number of PAX6+ cells at 6 weeks (F), total number of neurons at 13 weeks (G) and the fraction of PAX6+ cells at 13 weeks in culture. Total cell number, total number of PAX6 + cells and total number of neurons are all decreased in the CRISPR rescue-line compared to the parental patient line, confirming the causal effect of the single mutation on the observed phenotypes. Grey dots represent pooled control data points also displayed in figure 3A. Boxplots display median, first and third quartiles, and whiskers showing the largest and smallest values no further than 1.5 times the inter quartile range from first and third quartile, respectively. Data points outside of this range are plotted individually. * = p < 0.05; ** = p < 0.01; *** = p < 0.001

### Gene-ontology analysis based on RNAseq data corroborate abnormal NPC proliferation and neurogenesis processes

To evaluate the transcriptome changes associated with the phenotypes described above, we conducted bulk RNA sequencing (RNAseq) on 8-week old organoids. Principal component analysis (PCA) of the RNAseq data shows a clear separation in the transcriptional pattern between the two genotype groups, with 47% of the variance explain by principal component 1 (Fig 5A). Differential gene expression analysis identified a total of 339 differentially expressed genes (false discovery rate adjusted p-value < 0.05), including 89 genes that are upregulated and 250 genes that are downregulated in patient-derived organoids compared to controls (Fig 5B) (Table S3). To investigate whether these differentially expressed genes have known roles in biological processes related to the observed disease phenotypes, we performed Gene Ontology (GO) analysis (25) and found a statistically significant enrichment for genes involved in various biological processes, including cell proliferation and neurogenesis (Fig 5B, S4A and S4B and Table S4 and S5).

**Figure 5.**
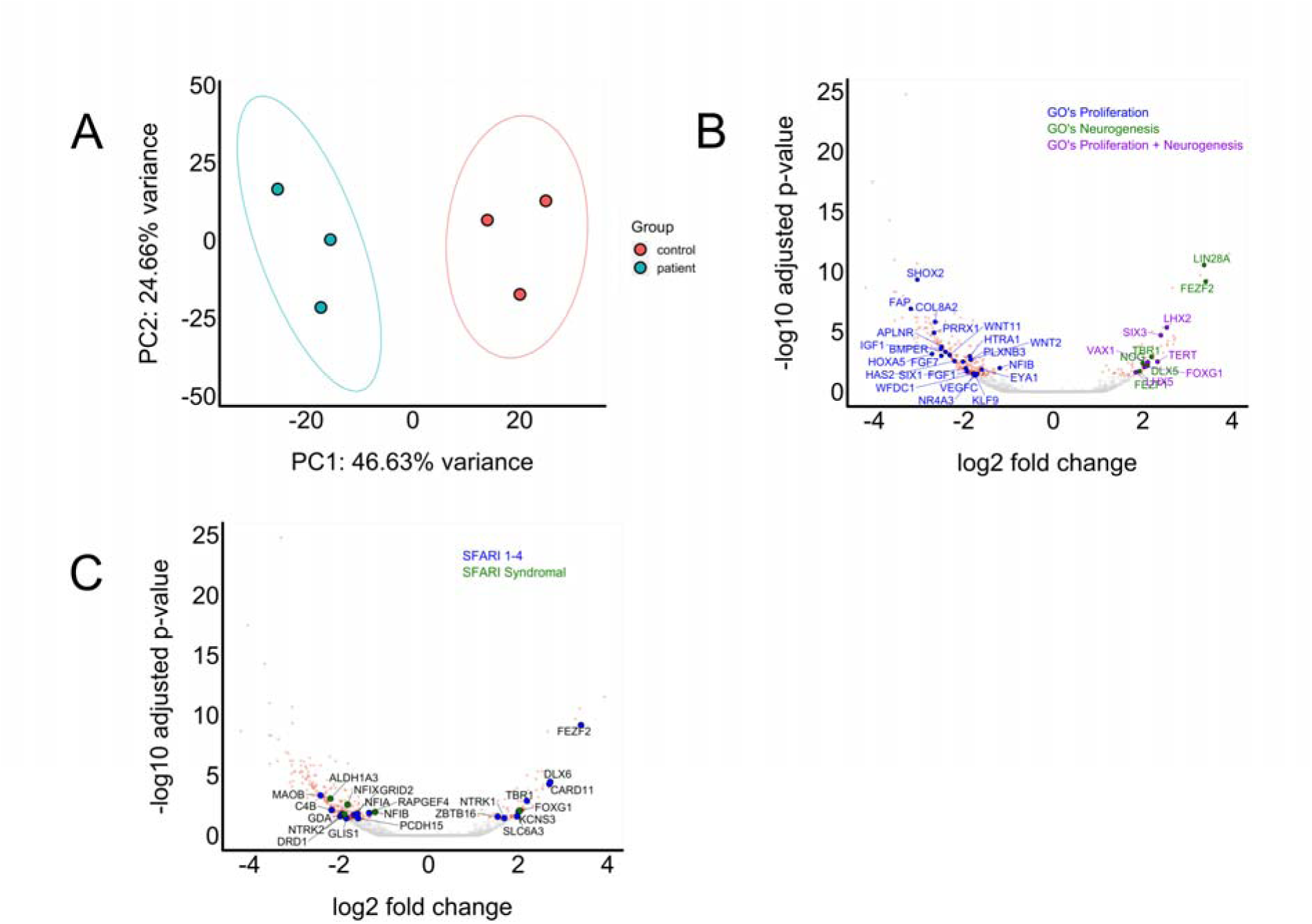
Differentially expressed genes are enriched for GOs related to cell proliferation and neurogenesis, as well as ASD-associated genes. **A.** Principal component analysis shows a clear distinction between the expression profiles of patient-compared to control-derived organoids at 8 weeks *in vitro*. **B.** Volcano plot showing differentially expressed genes between patient- and control-derived organoids at 8 weeks *in vitro*. Genes reaching statistical significance are colored in red, and genes below statistical significance threshold are colored in grey. A total of 250 genes are downregulated while 89 are upregulated. Genes are highlighted according to their Gene Ontology (GO). GOs related to cell proliferation (blue) (GO:0050673, GO:0050678, GO:0010463, GO:0010464, GO:0050679), neurogenesis (green) and both proliferation and neurogenesis (purple) (GO:2000177, GO:0072091, GO:0061351, GO:1902692, GO:0072089, GO:2000179, GO:0007405, GO:0050768, GO:005076). **C.** The same volcano plot that is shown in panel B, but now ASD-associated genes curated by SFARI are highlighted. Blue dots are genes in SFARI categories 1-4, green dots are in category ‘syndromic’.

### Differentially expressed genes show enrichment for ASD-associated genes and ASD Weighted Gene Co-expression Networks

Because ASD has been described as one of the core features of the neurodevelopmental syndrome associated with the homozygous c.3709DelG *CNTNAP2* mutation in the Old-Order Amish patients (7), we set out to investigate whether affected genes are enriched for genes with an established association with ASD. An enrichment could suggest that different genetic predispositions could lead to autism through effects on overlapping sets of genes. To do so, we compared the differentially expressed genes to the ASD-associated genes curated by SFARI Gene (31). We excluded genes in SFARI categories 5 and 6 as these were described as ‘hypothesized but untested’ and ‘evidence does not support a role’. Of the 877 genes remaining in SFARI categories 1-4 and category titled ‘syndromic’, 22 were differentially expressed between patient- and control-derived organoids (see table S6 for gene list). To test whether this enrichment reached the threshold for statistical significance, we performed permutation analyses (n = 10000). In each iteration, we generated a random gene list of equal length to the differentially expressed genes and found that this was indeed the case (z = 3.26, p=0.001) (Fig S4C).

We subsequently used the same approach to test whether the differentially expressed genes were also enriched in three independently created Weighted Gene Co-Expression Networks (WGCNAs) for ASD (32–34). These WGCNAs were constructed based on the correlation of gene-expression levels between genes, and led to the generation of gene co-expression modules that are associated with specific biological or disease processes (35). We found a statistically significant enrichment for all three WGCNAs (Fig. S4C-F), indicating an overlap in transcriptional signature between the differentially expressed genes between patient and control-derived organoids and other forms of ASD (Fig 5BCD). We then compared the differentially expressed genes to gene regulatory network data publically available for other neuropsychiatric conditions, such as Alzheimer’s disease (AD) (36) and Bipolar Disorder (BD) (37), and found no statistically significant enrichment for these WGCNAs. This implies that the enrichment of differentially expressed genes in WGCNAs is specific to ASD and related neurodevelopmental disorders.

## DISCUSSION

In this study, we utilized forebrain organoids generated from hiPSCs derived from patients carrying the homozygous c.3709DelG mutation in *CNTNAP2* and healthy controls to investigate the effects of this mutation on cortical embryonic development. We discovered increased proliferation in PAX6-positive NPCs, leading to an increase in the generation of cortical neuronal- and non-neuronal cells. This increase in number of cortical cells is at least partially responsible for a corresponding increase in overall organoid volume in patient-derived organoids. By repairing the pathogenic mutation using CRISPR-Cas9, we were able to rescue these cortical overgrowth phenotypes, thereby confirming a causative effect of the homozygous *LoF* mutation in the context of an identical genetic background. Abnormal proliferation and neurogenesis were corroborated by GO analysis of genes that were differentially expressed between patient- and control-derived organoids. Finally, we show that there is an enrichment of these differentially expressed genes for ASD-associated genes and WGCNAs associated with ASD and other neurodevelopmental disorders.

The main findings from this study confirm abnormal brain development in processes analogous to those occurring during the first trimester of embryonic development. With this, the findings support the prevailing hypothesis of an early embryonic origin of the pathophysiology of ASD (38). The strongest evidence for increased cortical neurogenesis in ASD comes from post-mortem data from children with ASD and increased head circumference, reporting increased neuronal numbers ranging from 12% to 108% (30). This increase in neuronal number could be related to the increased cortical thickening seen in ASD, that has also been observed on MRI of patients with *CNTNAP2*-associated ASD. More recently, other studies deploying hiPSC-derived neuronal culture techniques have also reported increased NPC proliferation and increased neurogenesis in patients with ASD. This includes both monogenic forms of ASD (29) and idiopathic ASD (5, 39) – in both 2D and 3D neuronal cultures. These findings could be causally related to two clinical phenomena observed in patients with ASD that may have at least a partial overlapping etiology: early brain overgrowth and an increase in head circumference or macrocephaly (defined as head circumference > 97th percentile). A recent systematic review and meta-analysis that analyzed 27 studies reported macrocephaly in 15.7% of ASD cases 3% in controls (40). The same study also reported an increase in total brain volume (defined as brain volume 2 standard deviations above the mean) in 9% of patients with ASD.

Of note, the overgrowth phenotype described in this study has not been recapitulated in *Cntnap2* null mice. One study quantified the volume of both medial prefrontal and somatosensory cortices, and found no difference between null- and wildtype mice (41). Thus, although mouse models can capture some of the clinical features of a human condition such as ASD, important aspects of such a disease may be missed since these may exclusively be present in humans. Given the fact that one of the defining features of the human brain compared to other mammals is its expanded gyrencephalic neocortex (42) that is thought to give rise to our abilities of higher order brain processes, such as cognition and language, it is not entirely surprising that a truncation of a protein with a critical role in cortical development could have different phenotypic outcomes in different mammalian species. Species-specific differences in the role that *CNTNAP2* plays in brain development is reflected in its differential expression pattern as well. This was shown using *in situ hybridization*, revealing that gene expression is more diffusely distributed throughout the entire brain in rodents while more focal in humans with a preferential localization in prefrontal cortex (28).

The findings from this study lead to interesting questions regarding the development of novel treatments for neurodevelopmental disorders, such as *CNTNAP2*-associated ASD. It is critical to determine the extent to which pathophysiological processes that occur as early as in the first trimester of pregnancy remain reversible after birth. Are the cortical overgrowth phenotypes critical to cognitive- and behavioral deficits seen in these patients, and if so, should treatments target these early developmental processes, and when? In this study, we show that the field of brain organoid modeling holds great promise for working towards a preclinical modeling system with face validity, with the ultimate goal of creating targeted treatments that may eventually improve the lives of patients with severe neurodevelopmental disorders such as *CNTNAP2*-associated ASD.

## Acknowledgements

The authors would like to thank Dr. Yuanjia Wang and Chen Chen for their advice on the statistical analyses using the linear mixed model. We thank Dr. Maura Boldrini for her advice on the BrdU labeling experiments and Yelizaveta Gribkova for performing Western blotting experiments. We thank Drs. Alexander Sosunov, Andrew Dwork and Gorazd Rosoklija for their advice on histology experiments. We thank Joseph Sall from NYU Langone’s Microscopy Laboratory (supported by the Cancer Center Support Grant P30CA016087 at the Laura and Isaac Perlmutter Cancer Center) for technical assistance with the light sheet microscopy image acquisition and Dr. Raju Tomer for his advice on the light sheet analysis. We thank Dr. Theresa Swayne and Dr. Laura Munteanu for their advice on image processing and analysis at the Confocal and Specialized Microscopy Shared Resource of the Herbert Irving Comprehensive Cancer Center at Columbia University, supported by NIH grant #P30 CA013696 (National Cancer Institute).

## Author Contributions

S.M. and B.X. conceived the research project, J.O.J., C.L., Y.S., K.S. and G.C. performed experiments and collected data, J.O.J., B.X., B.M., F.P. and G.C. analyzed data, B.C. reprogrammed patients’ samples, S.M., K.S. and K.B. were involved in the recruitment of the patients. S.M., B.X., C.K., J.A.J., S.K., J.O.J contributed to design of the research project and discussion of the data. J.O.J., S.M., B.X., J.A.J. and C.K. wrote the manuscript with input from the other authors.

## Competing interests

The authors declare no competing interests.

