## Supplemental table 1 for "Cortical Overgrowth in a Preclinical Forebrain Organoid Model of *CNTNAP2*-Associated Autism Spectrum Disorder"

| **Antibody** | **Vendor** | **Catalog** | **Species** | **Dilution IHC** | **Dilution WB** |
| --- | --- | --- | --- | --- | --- |
| MAP2 | Abcam | ab5392 | Chicken | 1:1000 |  |
| PAX6 | Biolegend | 901302 | Rabbit | 1:150, 1:20* |  |
| CNTNAP2 A | Genscript | A01426-100 | Rabbit |  | 1:1000 |
| CNTNAP2 B | Neuromab | K67/25 |  |  | 1:1000 |
| Ki67 | Abcam | ab6326 | Rabbit | 1:300 |  |
| NeuN | Millipore | ABN78C3 | Rabbit | 1:20* |  |
| BrdU | Abcam | ab6326 | Rat | 1:400 |  |

* Concentration used for isotropic fractionation experiments
