## Supplemental table 2 for "Cortical Overgrowth in a Preclinical Forebrain Organoid Model of *CNTNAP2*-Associated Autism Spectrum Disorder"

| Donor template flanking c.3907DelG mutation | GGCGAGCTGGTGGAGTCCAACTGCGGGGCCTCGCCGCTGACCCTCTCCCCCATGTCGTCCGCCACCGACCCCTGGCACCTGGATCACCTGGATTCAGGTAAAGTCTTCAGCAACCTCAGGCAGGTTGCTTCATTTCTTTAAACCTCAGTCTCTTGGGACTATGGAATGGATTTAATAACAG |
| --- | --- |
| >CNT_T1-mut fwd gRNA oligonucleotide | CACCGGGTCGGTGGCGGACGACATG |
| >CNT_T1-mut rev gRNA oligonucleotide | AAACCATGTCGTCCGCCACCGACCC |
| > CNT_T11-mut fwd gRNA oligonucleotide | CACCGGCACCTGGATCACCTGATTC |
| > CNT_T11-mut rev gRNA oligonucleotide | AAACGAATCAGGTGATCCAGGTGCC |
| *CNTNAP2* c.3907DelG genotyping primer fwd | ACAACACCCCAGGATTCACT |
| *CNTNAP2* c.3907DelG genotyping primer rev | TCCTTTCATTTGAGCCCTGT |
